# Recapitulating the length polymorphism of RS3, a composite microsatellite implicated in human social behavior, via recombination in *Saccharomyces cerevisiae*

**DOI:** 10.1101/2025.11.09.687482

**Authors:** Catie E. Kean, Elina A. Radchenko, Noah M. Lindenbaum, Grace Materne, Sergei M. Mirkin

## Abstract

Arginine Vasopressin (AVP) through its receptor V1a, pivotally modulates social and sexual behaviors in mammals. Differences in gene expression and the resulting behavioral changes have been linked to variation in the 5’ flanking region of the encoding gene *AVPR1A.* In particular, length polymorphism in the composite microsatellite RS3 ((CT)_4_TT(CT)_n_(GT)_m_) is responsible for multiple alleles of *AVPR1A* in the human population. However, the source of this length variation is unknown. We established a genetically tractable experimental system in *Saccharomyces cerevisiae* to study the mechanism of instability in the RS3 microsatellite. We successfully recapitulated the full spectrum of the RS3 length polymorphism observed in humans and found that it results from genetic recombination between RS3 sequences. Ultimately, we conclude that R-loop associated genome instability elevates recombination rate and triggers repair via single strand annealing which can result in microsatellite length polymorphism. We speculate that recombination is responsible for RS3 length polymorphism in humans.

## Introduction

Short tandem repeats (STRs), otherwise known as microsatellites, consist of 1-to-9 bp repetitive units (Liu et al., 2020). These sequences comprise 1-3% of the human genome and are associated with heightened instability (Lander et al., 2001). Most notably, massive expansions of STRs result in over 60 hereditary diseases, collectively termed repeat expansion disorders (REDs) (reviewed in Chen et al., 2025). However, even small variations in repeat length have been shown to alter gene expression by modifying chromatin structure, transcription, or splicing (Bagshaw, 2017).

Beyond pathogenicity, length polymorphism of microsatellites has been implicated in differences in social and sexual behavior (Hopkins et al., 2012). A notable example comes from prairie voles (*Microtus ochrogaster*) and meadow voles (*Microtus montanus*). While these species are nearly identical, they exhibit distinct social behaviors. Meadow voles are asocial, non-parental, and promiscuous, while prairie voles are highly affiliative, biparental, and monogamous (Young et al., 1999). This difference is attributed to decreased expression of the arginine vasopressin receptor V1a (V1aR) in the ventral forebrain of meadow voles relative to prairie voles (Hammock and Young, 2004). For example, when an AAV vector carrying the prairie vole *AVPR1A* gene was injected into the ventral pallidum of male meadow voles, they exhibited an increase in receptor density and monogamous behavior (Lim et al., 2004). In contrast, when *AVPR1A* expression in the ventral pallidum of prairie voles was inhibited via shRNA, males with low V1aR densities displayed impaired partner preference (Barrett et al., 2013).

*AVPR1A* is 99% homologous between the two vole species, the difference lies in a region approximately 760 bp upstream of the transcription start site. Prairie voles possess a complex microsatellite region consisting of di- and tetranucleotide STRs, which is nearly absent in meadow voles (Young et al., 1999). This microsatellite region appears to upregulate expression of *AVPR1A* and alter receptor distribution, accounting for behavioral divergence between the two species (Hammock and Young, 2004).

Human *AVPR1A* is highly similar to the corresponding gene in voles, however, the composition of the microsatellite repeats in the region upstream of the promoter differs significantly. A (GT)_n_ repeat resides at the −3956 position, followed by a complex microsatellite, RS3 ((CT)_4_TT(CT)_n_(GT)_m_) at −3625, and a (GATA)_n_ microsatellite, RS1, at −500. As with voles, variation in the length of the RS3 microsatellite has been linked to significant divergences in mating, aggression, and sexual behaviors (Young and Donaldson 2013; Sadino and Donaldson 2018). There are at least 16 distinct alleles of *AVPR1A* due to RS3 length polymorphism (Thibonnier et al., 2000; Erasmus et al., 2015). Alleles with shorter RS3 repeats have been linked to social anxiety, poor pair bonding, autism, and aggression in humans (Zhang et al., 2020; Meyer-Lindenberg et al., 2009; Thompson et al., 2024; Walum et al., 2008; Procyshyn et al., 2017; Vollebregt et al., 2021). Thus, length polymorphism in the RS3 microsatellite produces a diverse array of biologically relevant phenomena. However, the molecular mechanisms responsible for this variability remain elusive.

The RS3 sequence possesses the intrinsic capability to form alternative DNA structures. The (GT)_n_ component could form left-handed Z-DNA, while its (CT)_n_ repeats could form triplex H-DNA (Khristich & Mirkin 2020). Numerous studies have demonstrated that these alternative structures perturb DNA replication and repair, resulting in expansions and contractions, a common cause of RED phenotypes (Li et al., 2025). They can also lead to the formation of double-stranded breaks in DNA, stimulate homologous recombination (Khristich and Mirkin, 2020), trigger gross-chromosomal rearrangements (McGinty et al., 2017), and elevate mutagenesis both locally and at a distance (Shah and Mirkin 2015). Thus, the structural features of the RS3 microsatellite could be at heart of its length variability. Alternatively, misalignment of microsatellites during recombination could account for the length variation. The latter mechanism has been documented in other polymorphic microsatellites (Appelgren et al., 1997).

Herein, we investigated the mechanisms responsible for RS3 length polymorphism with a genetically tractable yeast experimental system. We determined that individual RS3 repeats are stable and do not readily undergo expansions or contractions in the (GT)_n_ or (CT)_n_ components or trigger fragility. However, misalignment during intrachromosomal recombination between two RS3 repeats produces the full spectrum of variability present in the human population. Using candidate gene analysis, we found that R-loop formation during transcription through the repeat-containing cassette promotes the recombination event and produces sequence variation through misalignment during single-strand annealing (SSA).

## Results

### The RS3 microsatellite does not expand or promote fragility in yeast

We previously showed that structure-prone DNA repeats can expand and contract in yeast (Kim and Mirkin et al., 2013). Given that the RS3 sequence could be structure-prone, we first sought to determine if it possessed the ability to expand using our modified intronic yeast system (Shah et al., 2012; Shah et al., 2014; Shishkin et al., 2009). The RS3 microsatellite (CT)_4_TT(CT)_8_(GT)_24_ was inserted into an artificial *URA3* intron in the *P_GAL1_-UR-intron-A3* cassette generating the *P_GAL1_-UR-RS3-A3* construct (Fig. 1A). In total, the intron measured 973 bp. We also created a control cassette that contained a non-repetitive sequence of the same length as the RS3 repeat. Using a *TRP1* selection marker, these constructs were, respectively, integrated into the yeast genome 1 kb downstream of the ARS306 replication origin. Since yeast lack the ability to splice out introns greater than approximately 1 kb, expansions of the RS3 that add 40-50 base pairs were expected to inactivate the reporter’s splicing, making yeast resistance to 5-FOA.

**Figure 1:**
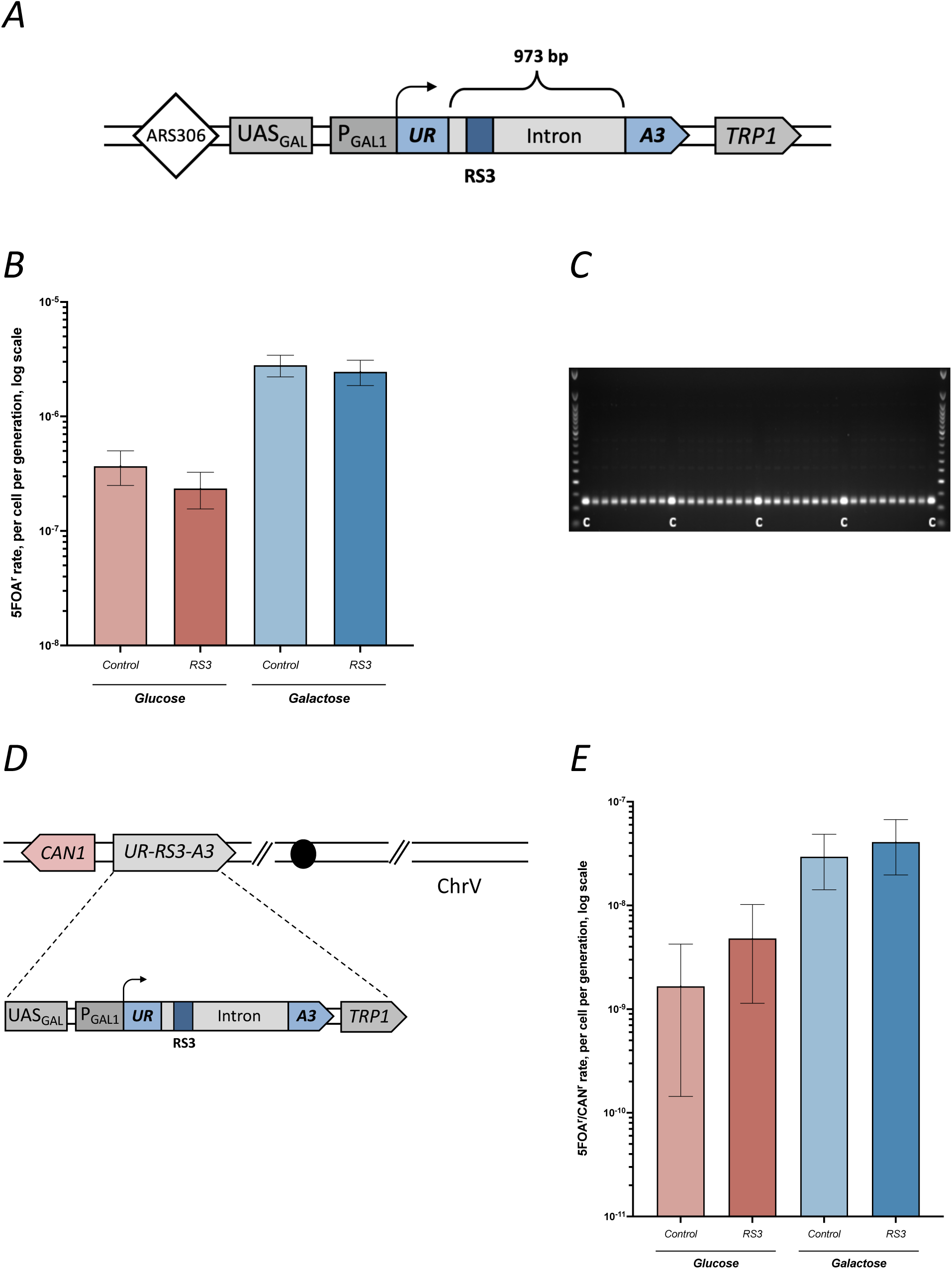
A system to detect expansions in the RS3 repeat. (A) The RS3 sequence was inserted into a 973 bp intron. Large expansion of the RS3 region would result in *URA3* gene inactivation and 5-FOA resistance. (B) Expansion rates for strains harboring the construct depicted in (A) and “control” strains with non-repetitive sequence in place of RS3. Assays were conducted on both glucose and galactose. Error bars mark 95% confidence intervals. (C) PCR products demonstrating the lengths of the RS3 repeat tracts from colonies grown on 5-FOA plates. The non-expanded amplified region is expected to be 117 bp. A 50 bp ladder was used in the distal lanes. The control “C” lanes are amplified from the original construct with unexpanded RS3 repeats. (D) The *PGAL_1_-UR-RS3-A3* cassette from (A) was integrated upstream of the *CAN1* gene on the nonessential arm of chromosome V for an arm loss assay. The formation of a double stranded break leads to loss of the cassette and *CAN1* and thus resistance to 5-FOA and canavanine. (E) Expansion rates for strains harboring the construct depicted in (D) and “control” strains with non-repetitive sequence in place of RS3. Assays were conducted on both glucose and galactose. Error bars mark 95% confidence intervals.

Fluctuation assays, as described in Radchenko et al. (2018) were used to determine expansion rate. Individual colonies were first grown on non-selective complete media containing either glucose or galactose and supplemented with uracil, followed by plating on the selective plates containing galactose and 5-FOA. The 5-FOA^r^ phenotype alone does not confirm the presence of an expanded repeat, as *URA3* inactivation can also result from mutagenesis independent of microsatellite length changes. Thus, expansions must be confirmed by PCR of 5-FOA-resistant colonies. When no expansions occur, 5-FOA^r^ rate is equivalent to the rate of *URA3* mutagenesis. We observed no significant differences between the 5-FOA^r^ rates in strains harboring the RS3 construct and the non-repetitive control regardless of transcription status (glucose repression versus galactose induction) (Fig. 1B). Furthermore, PCR amplification and gel electrophoresis of RS3 repeats in 50 colonies taken from 5-FOA plates revealed no expansions (Fig. 1C). Thus, we concluded that RS3 does not readily expand in our intronic system.

We previously demonstrated that repetitive DNA regions and secondary structures can increase chromosome fragility (Kim et al., 2008; McGinty et al., 2017). Therefore, we wanted to examine whether the RS3 microsatellite triggers fragility using Kolodner’s arm loss assay as described in McGinty et al. (2017). The *P_GAL1_-UR-RS3-A3* cassette from the expansion assay was integrated centromere-proximal to the *CAN1* gene on the non-essential left arm of chromosome V (Fig. 1D). The cassette containing a nonrepetitive sequence in place of RS3 was integrated in a separate strain.

In the event of DSB formation, the left arm of chromosome V, including part of the *P_GAL1_-UR-RS3-A3* cassette and *CAN1* gene, would be lost. These events could be detected by colony growth on plates containing both canavanine and 5-FOA. As simultaneous *can1* and *ura3* mutations are extremely rare, colony growth on canavanine+5-FOA media was considered a proxy of arm loss mediated by a DSB. We performed arm loss assays with the control and RS3 constructs and with growth on both glucose and galactose containing media. We observed no significant differences in the rate of arm loss between the control sequence and RS3 on both glucose and galactose (Fig. 1E). Notably, however, transcription through these cassettes, induced by galactose, caused a 8.5-fold increase in arm loss for the RS3-cassette and a 17-fold increase for the non-repetitive control.

### RS3 microsatellites can recombine and generate length variation

To explore the propensity of the RS3 microsatellite to recombine, we created a new yeast experimental system to measure the rate of intrachromosomal recombination. In the *P_GAL1_*-*UR-RS3-ADE2-RS3-A3* cassette, two identical RS3 microsatellites of the base composition (CT)_4_TT(CT)_8_(GT)_24_ were inserted into the artificial intron of *URA3,* flanking the *ADE2* gene, and placed under the control of the galactose promoter (Fig. 2A). Before recombination, this composite intron measured 2824 bp, which is above the splicing length limit for yeast. Thus, cells carrying the *P_GAL1_*-*UR-RS3-ADE2-RS3-A3* cassette are Ura^−^in the presence of galactose while Ade^+^, since the intron contains a full copy of the *ADE2* gene (Hooks et al., 2014). Crucially, the two RS3 repeats are the only homologous sequences within this cassette. Recombination excises the *ADE2* gene, shortening the intron to 512 bp, thus restoring *URA3* gene function. These recombination events are, therefore, detected as Ura^+^ Ade^−^(red) colonies growing on galactose-containing media.

**Figure 2:**
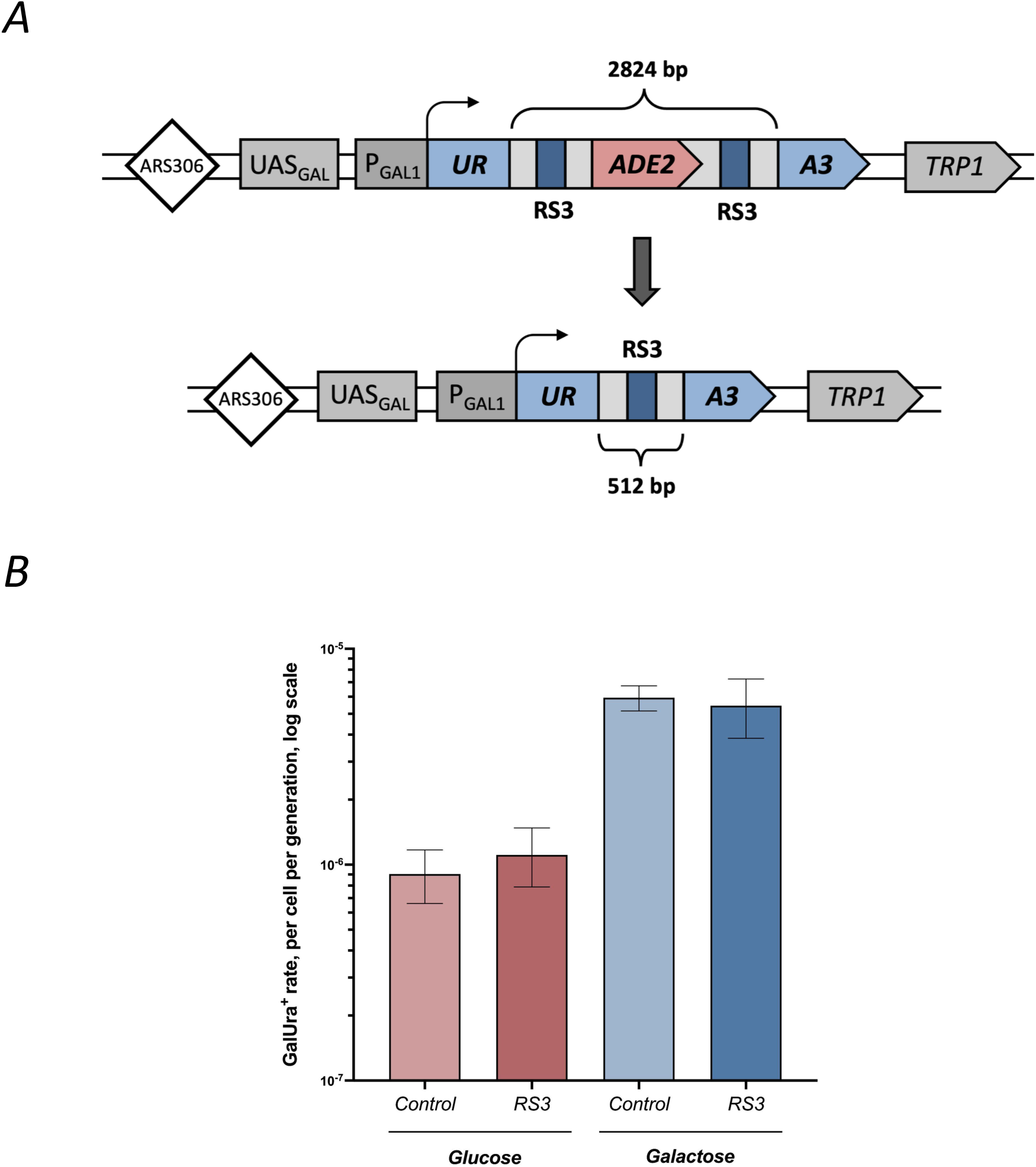
A system to detect recombination between RS3 repeats. (A) Two RS3 repeats, (CT)_4_TT(CT)_8_(GT)_24_, flanking *ADE2* gene were artificially inserted into an intron of the *URA3* gene under the *GAL1* promoter control. The resultant cassette – *UR-RS3-ADE2-RS3-A3 –* was integrated into the chromosome III adjacent to the *ARS306* replication origin. The total length of the intron is 2824 bp. Recombination events between the RS3 sequences shorten the intron to 512 bp, thus enabling splicing and excising the *ADE2* gene. (B) Recombination rates for strains harboring the construct depicted in (A) (blue) and strains with an identical construct, but in the place of the RS3 microsatellite a non-repetitive control sequence of the same length was integrated (red). Lighter columns represent cells grown on glucose during the nonselective phase and dark columns represent cells grown on galactose. Error bars indicate the 95% confidence intervals of each strain.

Fluctuation assays were used to calculate the recombination rate. Individual yeast colonies were first grown on the non-selective media containing either glucose or galactose, followed by plating onto the Ura^−^ selective media containing galactose to induce transcription. RS3 microsatellites recombined at a rate of ∼1.1 × 10^−6^ per cell per generation when the reporter was repressed in the presence of glucose (Fig. 2B). Transcriptional induction led to the 5-fold increase in recombination rate to ∼5.5 × 10^−6^ per cell per generation.

Rather unexpectedly, we observed similar recombination rates for the control cassette, which contained two identical non-repetitive DNA sequences of the same length as the RS3 sequence (Fig. 2B). Thus, we conclude that the RS3 repeat does not serve as a recombinational hot spot, at least in intrachromosomal recombination, regardless of transcriptional status.

We then investigated whether RS3 repeats could undergo length changes during recombination (Fig. 3A). As more recombination events occurred on galactose, we proceeded with these conditions. We extracted DNA from Ura+ Ade-colonies, amplified the region containing the RS3 repeat with PCR and visualized the products through gel electrophoresis. We observed both repeat expansions and contractions and determined that imperfect recombination occurs at a rate of ∼5.1 × 10^−7^ per cell per generation, which corresponds to ∼10% of all events. Subsequently, the spectrum of length polymorphism was constructed via Sanger sequencing (Fig. 3B). Variability was primarily present in the (GT)_24_ segment, with a minimum of 11 and maximum of 31 units (Fig. 3C). The (CT)_8_ region demonstrated a less pronounced range of 8 to 9 units. Thus, we were able to fully recapitulate length polymorphism of the GT segments of the RS3 repeat observed in humans in our yeast experimental system (Erasmus 2015).

**Figure 3:**
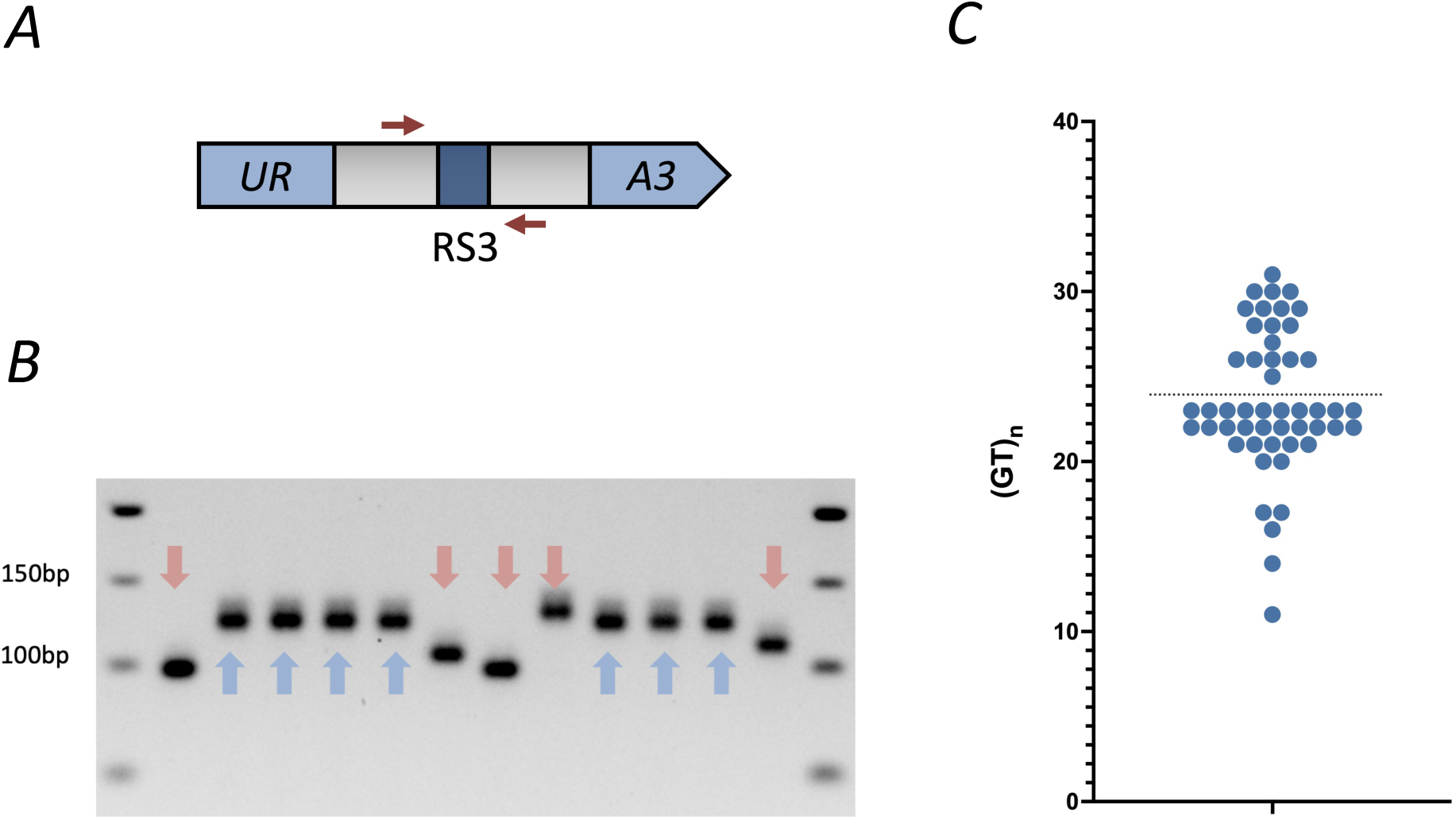
Length polymorphism in RS3 following recombination. (A) Schematic of primer annealing within *URA3.* Primers Opt1-2_in_s_fwd (top strand) and RS3_NgoMIV_short_R (bottom strand) were used to amplify the RS3 region in Ura+ colonies. (B) PCR products showing the lengths of the RS3 repeat tracts from colonies grown on selective plates. The amplified region is expected to be 120 bp, a 50 bp ladder is used in the distal lanes. Red arrows indicate “out-of-register,” or “imperfect” recombination events. Blue arrows show the target allele (CT)_4_TT(CT)_8_(GT)_24_, or “perfect” recombination events. (C) Length distribution of the (GT)_24_ repeat among 50 Ura+ Ade-colonies grown on selective plates. Colonies displayed polymorphism according to preliminary gel electrophoresis analysis.

### The genetic mechanism of RS3 recombination

The RS3 repeat in our recombination system measures 74 bp: (CT)_4_TT(CT)_8_(GT)_24_. This length is at the lower end for synthesis-dependent strand annealing (SDSA) and double-strand break repair (DSBR) — two key homologous recombination pathways in yeast (Pâques et al., 1999). Similarly, it satisfies the minimal length requirement for single-strand annealing (SSA) pathway but lies significantly below the optimal length. At the same time, it is much longer than the 5-to-20 bp microhomology required for microhomology-mediated end joining (MMEJ) pathway (McVey and Lee 2008). Therefore, it was not clear which pathway promotes recombination between RS3 repeats in our system. We utilized candidate gene analysis to address this question.

First, we assessed the involvement of key homologous recombination (HR) regulators in *S. cerevisiae*. Rad52 is a central effector of recombination in yeast and responsible for annealing homologous sequences; Rad59 is a Rad52 paralog and can partially compensate when Rad52 is lost (Mortensen et al., 1996; Shinohara et al., 1998; Sugawara and Haber, 1992; Sugawara et al., 2000). Indeed, loss of Rad52, Rad59, and both genes in tandem resulted in a ∼3-fold reduction in the recombination rate (Fig. 4). Strikingly, loss of Rad51 resulted in hyper-recombination in our system. Rad51 forms nucleoprotein filaments on single-stranded DNA and catalyzes homologous DNA pairing and strand exchange (Ogawa et al., 1993; Sung, 1994). Furthermore, loss of Rad54, which stabilizes Rad51 nucleoprotein filaments and promotes DNA strand exchange (Wright et al., 2014), also resulted in hyper-recombination. These results strongly argue against the involvement of SDSA or DSBR as both pathways require Rad51-mediated strand invasion. Instead, this data points to the SSA pathway, which depends on Rad52 and/or Rad59 (Symington et al., 2014).

**Figure 4:**
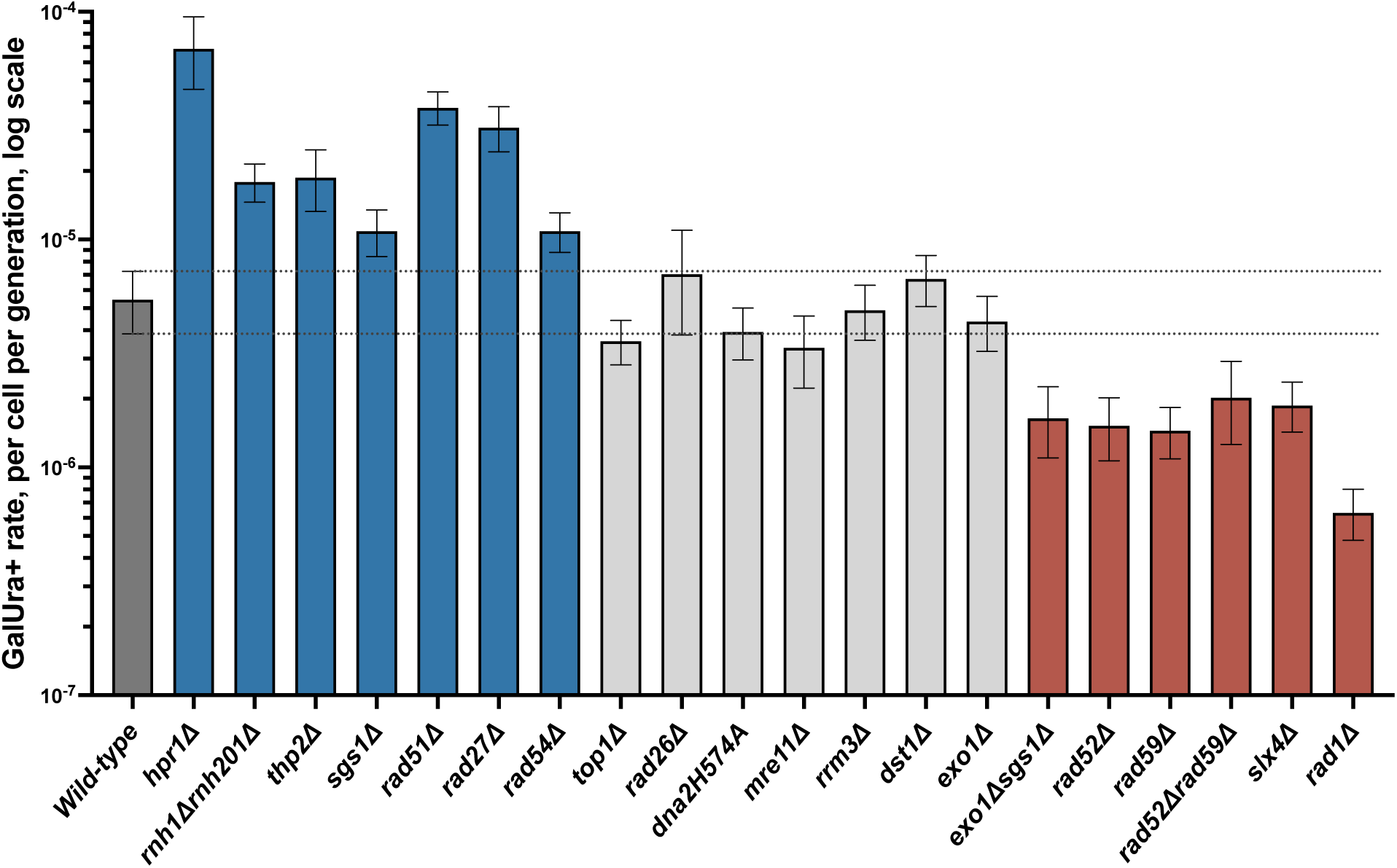
Genetic control of RS3 recombination. Error bars indicate the 95% confidence intervals of each strain. Dashed lines indicate the 95% confidence intervals of the wild-type strain.

In SSA, repetitive segments on either side of a double stranded break are annealed following resection (Fishman-Lobell et al., 1992). This process generates single-stranded flaps containing the sequences between repetitive regions. These flaps are then removed, and fill-in synthesis performed. The structure-specific endonuclease heterodimer Rad1-Rad10 is essential for the flap removal (Fishman-Lobell and Haber, 1992; Ivanov and Haber, 1995; Mazon et al., 2012). In agreement with the SSA hypothesis, we observed a substantial 10-fold decrease in recombination in *rad1Δ* mutants (Fig. 4). Furthermore, Slx4, a scaffold protein responsible for recruiting and regulating Rad1-Rad10 also showcased involvement (Flott et al., 2007). Strains deficient in Slx4 diplayed a 3-fold decrease in recombination.

Next, we examined the influence of end-resection proteins that generate single stranded overhangs. Mre11 is a component of the MRX complex (Mre11-Rad50-Xrs2) which is essential for the initial phase of DNA end resection during double-strand break repair (Mimitou and Symington, 2008; Zhu et al., 2008; Shim et al., 2010). Long range resection is facilitated by two independent ways: Exo1, a 5’→3’ exonuclease or Sgs1-Dna2 helicase-nuclease complex (Zhu et al., 2008). *MRE11* and *EXO1* gene knockouts did not affect recombination in our system (Fig. 4). Since *DNA2* is an essential gene, we created a *dna2-H547A* mutant, which possesses reduced endonuclease activity and impaired end-resection (Park et al., 2021). This mutation had no effect on RS3 recombination. However, *exo1Δsgs1Δ* double mutant showed a significant ∼3 fold decrease (Fig. 4) pointing to the role of the Sgs1-Dna2 complex in end-resection in our system. Intriguingly, the *sgs1Δ* mutant alone exhibited increased incidence of recombination. In addition to its roles in resection, Sgs1 can also resolve co-transcriptional R-loops which are known to promote the incidence of genetic recombination (Chang et al., 2017; Aguilera and García-Muse, 2012). Thus, while the *exo1Δsgs1Δ* double mutant is defective in resection, the full extent of this effect could be masked by an increase in recombination rate due to R-loop accumulation. Similarly, larger increases in recombination due to increased prominence of R-loops could be masked by Sgs1 mediated defects in resection.

This consideration warranted a closer look at the influence of R-loops in our system. DNA damage generated during transcription is often repaired by recombination (Aguilera, 2002). Thus, transcription-associated recombination (TAR) describes a pattern of conditions in which actively transcribed regions are commonly associated with recombinogenic repair (Gottipati and Helleday, 2009). Several mechanisms have been proposed to explain how transcription enhances recombination, including increased exposure of DNA to recombination proteins and DNA-damaging agents, collisions between replication machinery and the transcription apparatus or R-loops, as well as changes in DNA topology (Aguilera, 2002).

Therefore, we subsequently examined the role of proteins commonly implicated in TAR in our system (Aguilera, 2002). The THO complex counteracts TAR by ensuring proper messenger ribonucleoprotein (mRNP) synthesis and export (Chávez et al., 2000). This complex consists of four core proteins in yeast: Hpr1, Tho2, Mft1, and Thp2. Both Thp2 and Hpr1 are recruited specifically to transcriptionally active chromatin, where they function at the interface between transcription elongation and mRNA export (Chavez and Aguilera, 1997; Chávez et al., 2000; García-Rubio et al., 2008). Mutations in either gene result in R-loop accumulation and transcription-dependent hyperrecombination (Chavez and Aguilera, 1997). These THO proteins play a large role in our system, as deletions of *THP2* and *HPR1* genes result in hyperrecombination (Fig. 4).

To further elucidate the involvement of R-loops, we created strains deficient in both RNase H1 and RNase H2. Both proteins cleave RNA-DNA hybrids but have distinct substrate preferences. Rnh1 requires at least four consecutive ribonucleotides to function effectively, while Rnh201 (the catalytic subunit of RNase H2) can process single ribonucleotides (Lockhart et al., 2019). The double *rnh1Δ rnh201Δ* mutant completely abolishes RNase H activity resulting in R-loop accumulation (Amon and Koshland 2016). This double mutant displayed hyper-recombination between RS3 repeats, supporting the involvement of R-loops in the pathway (Fig. 4). Furthermore, absence of the 5’ flap endonuclease, Rad27, which has documented roles in R-loop resolution, also resulted in hyperrecombination (Liu et al., 2023)

We also investigated other potential mechanisms of TAR at RS3 microsatellites. Topoisomerase I (Top1) resolves topological stress during transcription, preventing stalling and R-loop formation (Aguilera 2002). Deletion of *TOP1* showed no significant effect (Fig. 4). We also observed no effect from the deletion of *DST1,* which encodes the general transcription factor TFIIS, as well as the Rrm3 helicase (Clark et al., 1991; Ivessa et al., 2003).

In addition to SSA, the Rad1-Rad10 complex also has a role in TAR owing to its involvement in nucleotide excision repair (NER) (Petermann et al., 2022). Transcription-coupled nucleotide excision repair (TC-NER) is a specialized DNA repair pathway that targets lesions on the actively transcribed DNA strand (Duan et al., 2020). Rad26 protein recognizes stalled polymerase complexes and recruits the repair machinery to remove the damage. We found no involvement of Rad26 and therefore suppose no involvement of TC-NER in our system (Fig. 4). Consequently, Rad1-Rad10 activity is likely confined to removal of 3’ tails during SSA and possibly, cleavage of the displaced DNA strand during R-loop formation.

## Discussion

The RS3 microsatellite in the human *AVPR1A* gene displays length variability across populations with major physiological, behavioral, and evolutionary implications. However, the mechanism(s) responsible for this length polymorphism remain unknown. We created a genetically tractable yeast experimental system to elucidate genetic pathways responsible for this phenomenon. The RS3 microsatellite consists of STRs that can form Z-DNA and H-DNA. Previous studies have shown that structure-prone DNA repeats undergo expansions (reviewed in Khristich & Mirkin 2020). In contrast, the RS3 microsatellite did not undergo expansions in our intronic reporter, as the sequence is likely significantly shorter than the threshold length for large-scale STR expansions detected by our system (72 bp versus 200 bp) (Fig. 1B). Additionally, using an arm-loss assay we found that the sequence does not induce fragility (Fig. 1E).

We then constructed a novel system to study intrachromosomal recombination between two RS3 microsatellites. While RS3 does not recombine at a greater rate than the non-repetitive control sequence (Fig. 2B), recombination results in frequent length changes, corresponding to 10% of events. This is likely due to misalignment of repetitive DNA strands during recombination (Fig. 3B). With our system we fully recapitulated the RS3 length polymorphism observed in the human population (Fig. 3C).

Recombination in our system is increased by active transcription of the cassette, pointing to the phenomenon of transcription associated recombination (TAR) (Aguilera et al., 2002). A central cog of this pathway is the accumulation of R-loops. Our data show that disruption of R-loop regulation at all levels, from prevention of aberrant elongation (THO complex), to unwinding (Sgs1), and finally, RNA cleavage (Rad27 and RNase H) (Liu et al., 2023) affects recombination between RS3 repeats (Fig. 4). Furthermore, we observed that transcription induction greatly increases chromosome fragility (Fig. 1E), supporting the general observation of transcription-mediated DSB formation.

The presence of R-loops is known to lead to the formation of DNA nicks (Petermann et al., 2022) primarily owing to the exposure of ssDNA on the non-template strand for transcription (Aguilera and García-Muse, 2012). For example, cytosine deaminases efficiently target ssDNA, converting cytosine to uracil, which triggers base excision repair, generating nicks (Lee et al., 2023). Additionally, structure specific nucleases that recognize R-loops introduce nicks (Su and Freudenreich 2017). Unrepaired nicks generated during co-transcriptional R-loops may become recombinogenic DSBs during DNA replication.

It was recently shown that DSBs arising from nicks are potent inducers of microsatellite instability (Li et al, 2024). We hypothesize, therefore, that transcription through our experimental cassette creates DSBs in between the two RS3 microsatellites; the SSA pathway then repairs the damage (Fig. 5). SSA depends on strand annealing by Rad52 and Rad59 proteins, and their absence strongly decreases recombination rate in our system. SSA also requires resection, which in our case, seems to depend on the Sgs1-Dna2 complex. Finally, Rad1-Rad10 nucleases are needed to remove flaps formed during SSA. We see that inactivation of this complex decreases recombination between RS3 repeats (Fig. 4). Furthermore, we found that improper alignment of RS3 microsatellites generates length variation. Notably, SSA has previously only been implicated in generating deletions between direct repeats and not polymorphism in intrinsic sequences (Bhargava et al., 2016).

**Figure 5:**
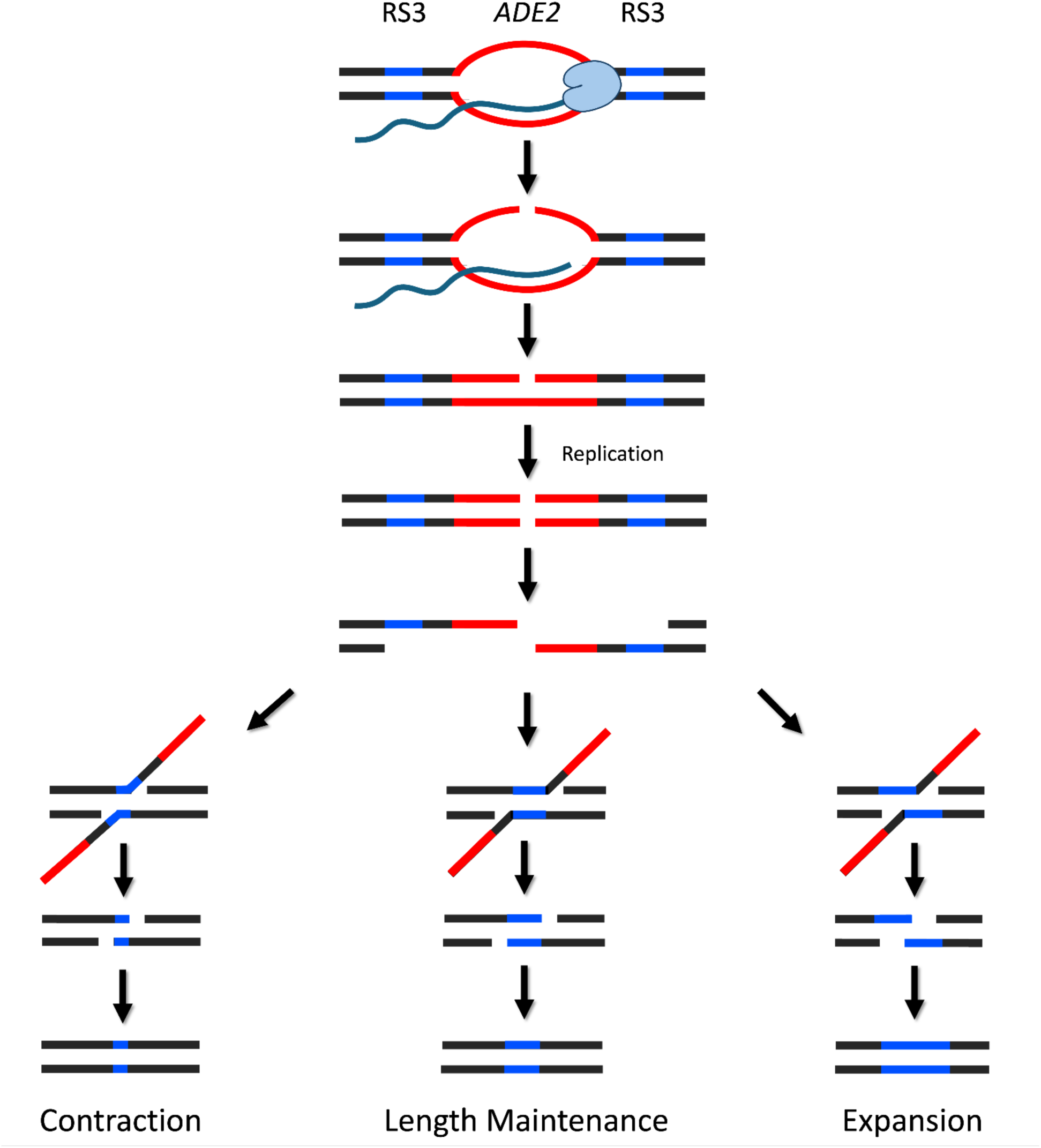
Transcription modulated genomic instability, either a DSB or a nick converted to a DSB during replication triggers SSA. Here, the damage is depicted in the *ADE2* locus. Exo1/Sgs1 perform resection to create single stranded RS3 homologies. Rad52 aligns the homologies and generates flaps. Rad1-Rad10 cleaves the 3’ flaps. Gaps are filled in and ligated and a single RS3 microsatellite remains. During a contraction event, out-of-register alignment of RS3 sequences results in partial cleavage of RS3 by Rad1-Rad10, shortening the repetitive region (left). Out-of-register annealing can also result in expansion events through incomplete alignment and fill in synthesis (right).

This model can also explain the inhibitory role of the Rad51 protein in our system (Fig. 4). When Rad51 is present, DSBs formed during DNA replication are preferentially repaired via sister chromatid exchange (SCE) rather than SSA. Repair via SCE would preserve the general structure of the cassette, *i.e. ADE2* gene flanked by RS3 repeats in the *URA*3 intron – an outcome that our system cannot detect.

Overall, we describe a general genetic event, in which ∼70 bp-long sequences on the same chromosome can recombine in a highly mutagenic and potentially detrimental manner. In our system, the alignment of these homologies results in a large deletion via TAR-mediated SSA and a currently uncharacterized transcription-independent pathway. We found that if short homologies are intrinsically repetitive, these processes produce high levels of length polymorphism. Furthermore, based on our data, this risk is compounded in actively transcribed regions, *i.e.* ∼60% of the human genome (ENCODE Project Consortium, 2012). The activity of this aberrant mechanism in coding or regulatory regions could result in deleterious health consequences. Future experiments are needed to test the limits of the recombination pathways through altering certain parameters such as the distance between homologous sequences, the length of homologous regions, and sequence identity.

## Methods

### Construction and integration of the RS3 expansion cassette

A *P_GAL1_-UR-RS3-A3* cassette was engineered to assess the RS3 microsatellite’s propensity to expand. First, the recombinant plasmid pYES3-G4G1C1-TET644 containing the *GAL1-UR-Intron-A3-TRP1* construct was generated by enzymatic digestion of two source plasmids (pYES3–G4G1C1 and PYES3-TET644) using NcoI and BglII, followed by fragment ligation. In this step, the *GAL1* promoter was obtained from pYES3-G4G1C1 and the *URA3* cassette was isolated from the pYES3-TET644 backbone vector. Successful assembly was verified through Sanger sequencing (Shah et al., 2012, 2014). Second, the RS3 sequence was synthesized with long self-dimerizing oligos Opt1-2_in_l_fwd and Opt2_in_l_rev, and then amplified with Opt1-2_in_s_fwd and Opt2_in_s_rev primers. The resulting RS3 fragment was inserted into the *URA3* intron of pYES3-G4G1C1-TET644 using restriction digest with NheI and PaeI, and ligation and subsequently transformed into *E. coli* Top10 cells. This digestion excised a 253 bp region and replaced it with the RS3 sequence. Additional cloning was performed to restore the initial 973bp total intron length. A 319 bp fragment was amplified from the pYES3-G4G1C1-TET644 plasmid using Opt3-PaeI-fwd and Opt3-rev primers. The PCR fragment and the pYES3-G4G1C1-TET644-RS3 plasmid with the shorter intron were digested with PaeI restriction enzyme, ligated together and then transformed into *E.coli* Top10 cells to get the final pYES3-G4G1C1-TET644-RS3 plasmid carrying the *P_GAL1_-UR-RS3-A3* cassette. Correct RS3 insertion and orientation were confirmed using PCR, restriction enzyme digestion, and Sanger sequencing, which matched the target allele (CT)_4_TT(CT)_8_(GT)_24_. The plasmids, pYES3-G4G1C1-TET644 (control) and pYES3-G4G1C1-TET644-RS3, were then digested with SmiI to integrate into the SMY706 yeast strain, creating YER1 and YER24 strains, respectively (Aksenova et al., 2013, Gietz and Schiestl, 2007). Successful transformation in yeast was confirmed by PCR. Proper integration and orientation were confirmed with Sanger sequencing.

### Measuring 5-FOA resistance

5-FOA^r^ rates were quantified using fluctuation assays based on the protocol developed by Radchenko et al. 2018. During the non-selective phase, individual colonies were grown on glucose-containing media (YPD) for three days, while colonies on galactose-containing media (YPGal) required four days of growth. In the selective phase, twelve colonies were suspended in sterile water and serial diluted by measures of 1:10 to achieve a maximum dilution factor of 10^5^. Following dilution, measured volumes from each colony suspension were plated on YPD (non-selective) media and on 0.1% 5-FOA media with galactose for selection. YPD plates were grown at 30 °C for three days and selective plates for five days before obtaining colony counts for both conditions. Colony counts from selective plates and corresponding non-selective plates were entered into the FluCalc program (Radchenko et al., 2018). This program employs the Ma-Sandri–Sarkar maximum likelihood estimation method. This method is considered one of the most faithful estimators of mutation rate as it compares practically infinite ranges of mutation rates and accounts for dilutions by incorporating plating efficiency calculations. 5-FOA^r^ rate differences were considered statistically significant when the 95% confidence intervals did not overlap between strains.

### Calculating expansion rate

Following the fluctuation assay, RS3 repeats were amplified from 5-FOA^r^ colonies and visualized through PCR and gel electrophoresis. Genomic DNA was extracted from randomly selected 5-FOA^r^ colonies using lyticase (Sigma-Aldrich L4025). The RS3 repeat was amplified using primers Opt1-2_in_s_fwd and Opt2_in_s_rev. PCR products were then visualized on a 1.5-2% agarose gel with 50 bp ladders (NEB #N0556S). Products were also amplified from the original RS3 plasmid, pYES3-G4G1C1-TET644-RS3, with no expansions to serve as controls.

### Construction and integration of the arm loss fragility constructs

To study arm-loss fragility events, two strains, YER5 and YER7, were created by integrating the expansion *P_GAL1_-UR-intron-A3* and *P_GAL1_-UR-RS3-A3* cassettes amplified with RM286-Cass_ChrVCAN1_intF and RM287-Cass_ChrVCAN1_int_R primers, on chromosome V proximal to a *CAN1* selection marker of the SMY923 strain (Aksenova et al., 2013).

### Calculating fragility rate

Both YER5 and YER7 strains were grown on YPD (6 days) or YPGal (7 days) to produce large colonies. Cultures were diluted and plated on YPD and 5-FOA/canavanine media with galactose. Selective plates enabled identification of arm-loss or gross chromosomal rearrangement events, as cells only survived when both *CAN1* and *URA3* were lost, conferring resistance to both drugs. During the selective phase YPD plates were incubated for 3 days, while selective plates were incubated for 5 days at 30°C. The frequency of 5-FOA/canavanine-resistant colonies was calculated as the rate of arm-loss (GCR, or Gross Chromosomal Rearrangements) events per cell per generation (Radchenko et al., 2018).

### Construction and integration of the RS3 recombination cassette

A *P_GAL1_-UR-RS3-ADE2-RS3-A3* cassette incorporated two RS3 repeats positioned within an artificial intron and an *ADE2* gene inserted between them. These RS3 regions represented the only homologous sequences within the entire construct. The system was assembled through stepwise cloning. The *ADE2* gene was amplified from yeast chromosome XV using ADE2-PaeI-F and ADE2-PaeI-R primers and inserted downstream of the first RS3 repeat of the pYES3-G4G1C1-TET644-RS3 plasmid using PaeI digestion and ligation. Correct insertion and orientation of *ADE2* was verified through PCR and sequencing. The second RS3 sequence was then synthesized with self-dimerizing oligos RS3-NgoMIV-long-F and RS3-NgoMIV-long-R, amplified with RS3-NgoMIV-short-F and RS3-NgoMIV-short-R and introduced into the intron downstream of *ADE2* using NgoMIV digestion and ligation. The completed cassette featured an intron spanning 2824 bp with RS3 sequences positioned 2238 bp apart. Following transformation of Top10 *E. coli*, proper orientation and length of the second RS3 sequence was verified through PCR and sequencing. An *ade2Δ* strain (YER9) was transformed with the cassette containing the dual RS3 repeats using a lithium acetate transformation protocol (Gietz et al., 2007), generating strain YER17. The cassette was positioned downstream of the ARS306 origin in a direct orientation, thus transcription and replication of the *UR-S3-ADE2-RS3-A3* cassette proceeded codirectionally.

### Calculating recombination rate

Recombination rates between RS3 microsatellites were quantified using a similar protocol to the expansion and arm-loss fragility assays. During the non-selective phase, individual colonies were grown on YPD for three days, while colonies on YPGal required four days of growth. For slow-growing strains (*hpr1Δ, dna2-H547A*), colonies were allowed to grow until reaching approximately 2mm in diameter. In the selective phase, YPD plates were incubated for three days at 30°C, and galactose-containing media lacking uracil (Gal/ura-) plates for five days. Recombination rate differences were considered statistically significant when the 95% confidence intervals did not overlap between strains.

### Repeat length analysis with single colony PCR

DNA was extracted from red colonies on selective media for PCR analysis. The recombined repeat was amplified using primers binding upstream of the first RS3 sequence (Opt1-2_in_s_fwd) and downstream of the second RS3 sequence (RS3_NgoMIV_short_R). PCR products were visualized using gel electrophoresis on 1.5-2% agarose gels until adequate separation was achieved. Bands were categorized as normal length, imperfect, or unclassified. Unclassified fragments underwent additional analysis on 2% agarose gels with wider wells. Control sequences from perfectly recombined RS3 fragments were loaded between unknown samples for comparison. The total frequency of imperfect recombination events was calculated for each strain. The proportions of ‘perfect’ and ‘variable’ recombination events were determined based on the random samples analyzed by PCR and then applied to the complete dataset. The exact sequence of imperfectly recombined RS3 repeats was assessed with Sanger sequencing to characterize polymorphism distribution.

### Gene deletion methodology

Candidate gene analysis was conducted through removing nonessential genes in *S. cerevisiae* from wild-type yeast strains YER17/1 or YER94/31 and employing either standard gene replacement techniques or CRISPR Cas9-mediated genome editing as described by Generoso et al. 2016. For conventional gene replacement without DSB induction, a plasmid containing a selectable marker (kanamycin (KAN), nourseothricin acetyl transferase (NAT), or hygromycin resistance (HYG)) was amplified using primers that incorporated homology to the targeted gene. For CRISPR-Cas9 knockout fragment construction, primers were designed with partially complementary sequences that were half homologous to a region within each target gene and half to the pRCC_N or pRCC_H plasmids (Generoso et al., 2016). The Cas9 plasmid backbone was amplified using Thermo Scientific high-fidelity Phusion polymerase with these target-specific primers producing linear DNA fragments encoding NAT- or HYG-resistance, Cas9, and the gene-targeting sgRNA. Repair template oligos were designed with overlapping sequences from the gene’s flanking regions and assembled via Phusion PCR. Specific information about knockout fragments, oligonucleotides used for synthesis, repair templates, and validation primers is available upon request.

Transformations were performed using a high-efficiency lithium acetate protocol modified from Gietz et al. 1992. Donor DNA was introduced into wild-type yeast using a solution containing 0.1M lithium acetate, 10 mg/ml salmon sperm DNA carrier, and 33% polyethylene glycol. For conventional gene replacement, 1000 ng of DNA knockout fragment was added, while for CRISPR-Cas9 editing, 2 μl of pRCC plasmid and 1000 ng of repair template were used. Transformation mixtures were incubated at room temperature for 25 minutes followed by 42°C for 25 minutes. Transformed cells were plated on YPD media supplemented with uracil and adenine (YPDUA) and replica-plated to selective media after 1-2 days, or grown in liquid YPDUA for one hour before direct plating on selective media. After two days, DNA was extracted from selective plate colonies and successful alteration was confirmed using locus specific primers. Fragment presence/absence and size were assessed using PCR and gel electrophoresis. Confirmed colonies were isolated on YPEG media and successful transformants stored at −80°C in liquid YPDUA with 14.2% glycerol.

## Notes

### Competing Interest Statement

The authors have declared no competing interest.

### Funding

- NIH: R35GM130322
- NSF: MCB 2153071

